## Supplementary material for "Causal reasoning over knowledge graphs leveraging drug-perturbed and disease-specific transcriptomic signatures for drug discovery"

This file contains:

- Supplementary Tables
- Supplementary Text
- Supplementary Figures

### Supplementary Tables

| Sr No. | Name | Reference | Description | Valid | Reason for not using |
|---|---|---|---|---|---|
| 1 | GEO | https://www.ebi.ac.uk/arrayexpress/ (from https://zenodo.org/record/4568170) | Disease | Yes | Potential disease dataset with Disease Ontology Identifiers (DOIDs) |
| 2 | Open Targets | https://www.opentargets.org/ | Disease | Yes | Potential disease dataset with Experimental Factor Ontology (EFO) identifiers |
| 3 | CREEDS | https://maayanlab.cloud/CREEDS/ | Disease / Chemical | Yes for drug only | Potential chemical dataset with pubchem compound identifiers |
| 4 | L1000 | https://lincsproject.org/LINCS/ | Chemical | Yes | Potential chemical dataset with pubchem compound identifiers |
| 5 | Broad | https://ctd2-data.nci.nih.gov/Public/Broad/ | Disease | No | Low number of chemicals (only 19) present |
| 6 | DFCI | https://ctd2-data.nci.nih.gov/Public/DFCI/ | Disease | No | Low number of diseases (only 12) present |
| 7 | HTI-PPI | https://ctd2-data.nci.nih.gov/Public/Emory | Chemical | No | High number of chemicals (2031) but each chemical has availability of 0-2 gene fold change present |
| 8 | SC2Diseases | http://easybioai.com/sc2disease/ | Disease | No | Low number of diseases (only 23) present |
| 9 | ToxicoDB | https://www.toxicodb.ca/datasets/ | Chemical | No | Only 10 chemicals are available |
| 10 | 10X Genomics | https://support.10xgenomics.com/ | - | No | Not available for download |
| 11 | Aging, Dementia and TBI study | http://aging.brain-map.org/download/index | Disease | No | Not relevant for our study |
| 12 | Allen Brain Atlas | http://human.brain-map.org/microarray/search | Disease | No | No gene expression data available for download |
| 13 | BioGPS | http://biogps.org/dataset/ | Disease | No | Multiple disease-specific datasets present |
| 14 | Cancer Genome Atlas (TCGA) | https://portal.gdc.cancer.gov/ | Disease | No | Multiple datasets with cancer-specific gene expression data |
| 15 | Cancer Therapeutic Response | https://ctd2-data.nci.nih.gov/Public/UTSW/Lung_Cancer_Natural_Products_Screening/ | Chemical | No | Multiple datasets present but no fold change data found |
| 16 | Cancerrxgene | https://www.cancerrxgene.org/downloads/anova | Chemical | No | No fold change |
| 17 | CellMiner | https://discover.nci.nih.gov/cellminer/home.do | Chemical | No | No transcriptomic data available |
| 18 | DeepCE | https://github.com/pth1993/DeepCE | Chemical | No | Subset of L1000 dataset |
| 19 | DeepMap | https://depmap.org/portal/download/ | Disease | No | No disease/drug fold changes found |
| 20 | GDA | http://gda.unimore.it/index.php | Drug | No | Not available for download |
| 21 | GENT2 | http://gent2.appex.kr/gent2/ | Disease | No | Tissue related gene fold data |
| 22 | HNC | http://hncdb.canceromics.org/ConnectiveMap | Chemical | No | Not available for download |
| 23 | MARINa | https://ctd2-data.nci.nih.gov/Public/Columbia/Neuroblastoma_TEAD4_MYCN/ | - | No | Gene fold change specific to neuroblastoma |
| 24 | MGI | http://www.informatics.jax.org/expression.shtml | - | No | Mouse related gene expression data |
| 25 | PulmonoDb | http://pulmondb.liigh.unam.mx/index.html | Disease | No | COPD specific dataset |
| 26 | Recount | https://jhubiostatistics.shinyapps.io/recount/ | - | No | Not relevant for our study |
| 27 | Transcriptional data | https://doi.org/10.1016/j.drudis.2012.07.014 | Disease/ Chemical | No | No dataset mentioned |

**Supplementary Table 1. Investigated datasets.**

| Sr No. | Name | Number of diseases/drugs mapped to any of the KGs | Average number of genes measured | Minimum number of genes measured | Maximum number of genes measured |
|---|---|---|---|---|---|
| 1 | GEO | 18 | 6569.55 | 26 | 18,123 |
| 2 | Open Targets | 44 | 958.57 | 1 | 3,072 |
| 3 | CREEDS | 39 | 556.97 | 539 | 570 |
| 4 | L1000 | 269 | 342.10 | 3 | 2,753 |

**Supplementary Table 2. Statistics on the genes measured in the four transcriptomics datasets used.**

| Network | Dataset | Drug-Disease Pairs from ClinicalTrials.gov | Unique Drugs (PubChem compound identifiers) | Unique Diseases (MONDO identifiers) | Possible Combinations |
|---|---|---|---|---|---|
| **OpenBioLink KG** | L1000-GEO | 296 (15.66%) | 189 | 10 | 1,890 |
| | L1000-Open Target | 385 (11.32%) | 189 | 18 | 3,402 |
| | CREEDS-Open Target | 146 (26.16%) | 31 | 18 | 558 |
| | CREEDS-GEO | 114 (36.77%) | 31 | 10 | 310 |
| **Custom KG** | L1000-GEO | 432 (12.83%) | 198 | 17 | 3,366 |
| | L1000-Open Target | 713 (9.23%) | 198 | 39 | 7,722 |
| | CREEDS-Open Target | 285 (24.36%) | 30 | 39 | 1,170 |
| | CREEDS-GEO | 171 (33.53%) | 30 | 17 | 510 |

**Supplementary Table 3. Clinical trial information mapped to the OpenBioLink and custom KGs.**

| Property | OpenBioLink KG | Custom KG |
|---|---|---|
| # Nodes | 4,831 | 8,489 |
| # Edges | 47,214 | 52,142 |
| # Activatory Edges | 29,022 | 43,578 |
| # Inhibitory Edges | 12,477 | 8,045 |
| Average degree | 17.17 | 12.161 |
| Longest shortest path in the KG | 11 | 11 |

**Supplementary Table 4. Properties of the OpenBioLink and custom KGs.**

| Network | Dataset | % of drug-disease pairs with a path length of 3 (drug-protein-disease) | % of drug-disease pairs connected by a path | % pairs with concordant paths after applying Step 2 of the algorithm |
|---|---|---|---|---|
| **OpenBioLink KG** | L1000-GEO | 2.75 | 70.80 | 0.37 |
| | L1000-Open Target | 2.79 | 53.44 | 0.61 |
| | CREEDS-Open Target | 12.19 | 61.47 | 0.58 |
| | CREEDS-GEO | 10.32 | 81.94 | 0.79 |
| **Custom KG** | L1000-GEO | 0.80 | 87.28 | 0.10 |
| | L1000-Open Target | 0.93 | 86.65 | 0.06 |
| | CREEDS-Open Target | 0.68 | 91.45 | 0.09 |
| | CREEDS-GEO | 1.37 | 92.55 | 0.42 |

**Supplementary Table 5. Percentage of drug-disease pairs at different steps for every dataset combination and KG.**

| - | OpenBioLink KG | Custom KG |
|---|---|---|
| **Method** | Precision (TP/TP+FP) | Precision (TP/TP+FP) |
| **Shortest paths** | 20.37% (33/162) | 10.41% (40/384) |
| **Common neighbours** | 34.61% (36/104) | 31.76% (27/85) |
| **Cosine similarity** | 43.75% (7/16) | 38.90% (14/36) |
| **Jaccard index** | 43.75% (7/16) | 37.14% (13/35) |
| **Sorensen index** | 43.75% (7/16) | 37.14% (13/35) |
| **Hub promoted index** | 34.34% (34/99) | 37.20% (16/43) |
| **Hub depressed index** | 43.75% (7/16) | 35.21% (25/71) |
| **Leicht-Holme-Newman index** | 43.75% (7/16) | 40% (14/35) |
| **Preferential attachment** | 5% (1/20) | 9.30% (4/43) |
| **Adamic-Adar** | 33.33% (30/90) | 32.143% (18/53) |
| **Resource allocation index** | 33.33% (30/90) | 32.143% (18/53) |

**Supplementary Table 6. Evaluation of the 11 benchmark methods using precision as a metric**. None of the benchmarked methods achieve a precision greater than 50%. Furthermore, we would like to note that most of the drug-disease pairs prioritized by each of these methods are the same since they are based on network proximity. Thus, if a drug and a disease share a large number of nodes, they will consistently be prioritized by most of these methods.

### Supplementary Text

#### 1. Processing of transcriptomic datasets

For datasets that did not already provide fold changes, we conducted differential expression analysis using the Limma R package (https://bioconductor.org/packages/release/bioc/html/limma.html) as described by https://zenodo.org/record/4568170. DEGs were then filtered to include only those with an adjusted $p$-value < 0.05.

#### 2. Benchmarked methods

Below, we introduce the 11 benchmarked methods based on network-similarly presented by Abbas *et al*. (2021) and Zietz *et al*. (2020).

1. **Shortest paths:** This method prioritizes a drug to its closest disease based on their shortest path (e.g., drug X -> gene Y -> disease Z).

2. **Common Neighbours (CN):** This method assigns higher scores to two nodes with a high number of common neighbours. Let $\Gamma(i)$ represents vector or set that contains the neighbors of node $i$. This method finds the number of neighbours that intersect between the two nodes.

$$s_{ij} = |\Gamma(i) \cap \Gamma(j)|$$

3. **Salton index (Cosine similarity):** This method measures the cosine of the angle between the columns in the adjacency matrix of the graph. This calculation is similar to the common neighbours. $k_i$ is defined as the degree of node $i$, or the number of neighbours node for node $i$.

$$s_{ij} = \frac{|\Gamma(i) \cap \Gamma(j)|}{\sqrt{k_i * k_j}}$$

4. **Jaccard index:** Similar to the previous two, this method finds the proportion of common neighbours and total neighbours between two nodes $i$ and $j$.

$$s_{ij} = \frac{|\Gamma(i) \cap \Gamma(j)|}{|\Gamma(i) \cup \Gamma(j)|}$$

5. **Sorensen Index:** Similar to Jaccard index, this method measures the relative size of an intersection between two sets of neighbours. This method came to rise through ecological community data.

$$s_{ij} = \frac{2 * |\Gamma(i) \cap \Gamma(j)|}{k_i + k_j}$$

6. **Hub Promoted Index (HPI):** This method incorporates common neighbours but assigns higher scores to hub nodes (high-degree nodes) because the denominator of this index relies on the minimum of degrees for both nodes.

$$s_{ij}=\frac{|\Gamma(i)\cap\Gamma(j)|}{min(k_i,k_j)}$$

7. **Hub Depressed Index:** In contrast to HPI, this measure assigns lower scores to links that are adjacent to hubs. This is the case because we find the maximum of degrees for both nodes in the denominator.

$$s_{ij}=\frac{|\Gamma(i)\cap\Gamma(j)|}{max(k_i,k_j)}$$

8. **Leicht-Holme-Newman Index (LHN-I):** This method can be seen as a variant of the common neighbours method, as it assigns high scores to common neighbour nodes while penalizing with respect to the degree of each node.

$$s_{ij}=\frac{|\Gamma(i)\cap\Gamma(j)|}{k_i*k_j}$$

9. **Preferential Attachment (PA):** This method is established on the fact that nodes that have higher links will form more future connections. Therefore, this link prediction model is just the product of the degree of both nodes.

$$s_{ij}=k_i*k_j$$

10. **Adamic-Adar (AA):** This method is based on the assumption that less connected nodes should be given more weight for link prediction. This is accomplished through simple counting of common neighbours and assigning weights to the nodes inversely proportional to the logarithm of their respective degrees.

$$s_{ij}=\sum_{z\in\Gamma(i)\cap\Gamma(j)}\frac{1}{logk_z}$$

11. **Resource Allocation Index (RA):** Similar to the Adamic-Adar index and inspired by the resource allocation process, this method essentially measures how much resource is communicated between two nodes *i* and *j*.

$$s_{ij}=\sum_{z\in\Gamma(i)\cap\Gamma(j)}\frac{1}{k_z}$$

### Supplementary Figures

**Function 1** Algorithm to prioritize protein targets for a given disease by correlating disease-specific transcriptomic signatures.

```
function IS_DRUG_PRIORITIZED(KG, disease, lmax, errors_allowed)
    proteins ← GET_PROTEINS(KG)
    disease_tr ← GET_TRANSCRIPTOMICS(disease)

    prioritized_proteins ← ∅
    for all protein ∈ proteins do
        paths ← GET_ALL_PATHS(KG, protein, disease, lmax)
        for all path ∈ paths do
            if IS_CONCORDANT(KG, path, disease_tr, errors_allowed) then
                prioritized_proteins.insert(protein)
                break
            else
                path_comp ← path ∗ −1                ▷ check complementary path
                if IS_CONCORDANT(KG, path_comp, disease_tr, errors_allowed) then
                    prioritized_proteins.insert(protein)
                    break
                end if
            end if
        end for
    end for
    return prioritized_proteins
```

**Supplementary Figure 1. Pseudocode of the RPath algorithm designed for target prioritization.**
